# Assembly by Reduced Complexity (ARC): a hybrid approach for targeted assembly of homologous sequences

**DOI:** 10.1101/014662

**Authors:** Samuel S. Hunter, Robert T. Lyon, Brice A. J. Sarver, Kayla Hardwick, Larry J. Forney, Matthew L. Settles

## Abstract

Analysis of High-throughput sequencing (HTS) data is a difficult problem, especially in the context of non-model organisms where comparison of homologous sequences may be hindered by the lack of a close reference genome. Current mapping-based methods rely on the availability of a highly similar reference sequence, whereas *de novo* assemblies produce anonymous (unannotated) contigs that are not easily compared across samples. Here, we present Assembly by Reduced Complexity (ARC) a hybrid mapping and assembly approach for targeted assembly of homologous sequences. ARC is an open-source project (<u>http://ibest.github.io/ARC/</u>) implemented in the Python language and consists of the following stages: 1) align sequence reads to reference targets, 2) use alignment results to distribute reads into target specific bins, 3) perform assemblies for each bin (target) to produce contigs, and 4) replace previous reference targets with assembled contigs and iterate. We show that ARC is able to assemble high quality, unbiased mitochondrial genomes seeded from 11 progressively divergent references, and is able to assemble full mitochondrial genomes starting from short, poor quality ancient DNA reads. We also show ARC compares favorably to *de novo* assembly of a large exome capture dataset for CPU and memory requirements; assembling 7,627 individual targets across 55 samples, completing over 1.3 million assemblies in less than 78 hours, while using under 32 Gb of system memory. ARC breaks the assembly problem down into many smaller problems, solving the anonymous contig and poor scaling inherent in some *de novo* assembly methods and reference bias inherent in traditional read mapping.

## INTRODUCTION

High-throughput sequencing (HTS) techniques have become a standard method for producing genomic and transcriptomic information about an organism (Schbath et al. 2012). The Illumina, Roche, and Life Sciences sequencing platforms produce millions of short sequences referred to “reads” that range in length from 50 to 700 base pairs (bp) depending on chemistry and platform. In shotgun sequencing, these short reads are typically produced at random, making them effectively meaningless without further analysis. The primary challenge in the analysis of HTS data is to organize and summarize the massive number of short reads into a form that provides insight into the underlying biology. Two analysis strategies, *de novo* sequence assembly and sequence mapping have been widely adopted to achieve this end.

The objective of *de novo* assembly is to piece together shorter read sequences to form longer sequences known as contigs. Sequence assembly is a challenging problem that is made more difficult by characteristics of the sequenced genome (e.g., repeated elements and heterozygosity) and by sequencing technology characteristics (e.g., read length and sequencing errors). Additionally, assembly algorithms are computationally intensive for all but the smallest datasets, thus limiting their application (Li et al. 2012). Finally, *de novo* assembly of large datasets typically produces many short contigs that require additional organization and analysis. Despite many advances and a large selection of assembly software packages, fragmentation and misassembly remain common problems and improving the quality of *de novo* sequence assemblies continues to be an area of active research (Bradnam et al. 2013).

Sequence mapping is often the first step carried out in resequencing projects where a good reference sequence exists. The objective of mapping is to align short reads against a reference sequence, thereby permitting direct sequence comparisons between a sample and the reference. This approach is significantly faster than *de novo* sequence assembly and has proven to be very effective at identifying sequence variants at a large scale (The 1000 Genomes Project Consortium et al. 2010). Unfortunately, this approach is entirely dependent on a reference sequence that is similar to the organism being sequenced. Differences between a sample and reference sequence (e.g., structural variations (SVs), novel sequences, an incomplete or misassembled reference, or sequence divergence) can result in unmapped or poorly-mapped reads, which may result in false variant calls (Li, 2011). In the context of RNA-Seq experiments, unmapped reads result in counting errors, and can affect the identification of differentially expressed genes (Pyrkosz et al. 2013). Resequencing projects are performed to identify differences between a sample and an established reference; however, the regions that are most divergent can also be the most difficult to map reads against. Because of this, mapping based approaches are inherently biased by the reference and only provide reliable results when sequence divergence is below the threshold at which reads can be mapped accurately.

The two approaches described above (mapping and *de novo* assembly) have been developed and optimized for whole-genome analysis; however, another class of problems exists in which specific regions of a genome or subsets of the sequenced DNA are analyzed. This type of analysis is appropriate in many instances, including sequence capture, viral genome assembly from environmental samples, RNA-Seq, mitochondrial or chloroplast genome assembly, metagenomics, and more. In cases like these, it has been necessary to develop custom pipelines to carry out analyses. In order to assemble the mammoth mitochondrial genome from whole genome shotgun data, Gilbert et al. (2007) first mapped the reads to a reference mitochondrial sequence, filtered the mapped reads and then “assembled using scripts to run existing assembly software”. Other tools that have been developed to address sub genome assembly include: MITObim an extension of the MIRA assembler (Hahn et al. 2013; Chevreux et al. 1999). It requires the user to first perform a mapping based assembly with MIRA, then to use the output of this assembly to do iterative read recruitment and assembly; however, according to the documentation, MITObim does not take advantage of paired-end reads for recruitment or extension. Further, it is not optimized for multiple targets or multiple samples, due to many steps that are manually carried out. The Mapping Iterative Assembler (MIA) uses an iterative mapping and consensus calling approach https://github.com/udo-stenzel/mapping-iterative-assembler). The algorithm is tuned for ancient DNA and was reported by Hahn et al. (2013) to be very slow. It also appears to only function with a single sample and reference; Other groups such as Malé et al. (2014), and Picardi and Pesole (2012) have also developed strategies for assembling smaller subsets from larger datasets; however, none of these were developed as a general purpose, highly parallelized homologous sequence assembler.

To address this problem we introduce a hybrid strategy, Assembly by Reduced Complexity (ARC) that combines the strengths of mapping and *de novo* assembly approaches while minimizing their weaknesses. This approach is designed for the myriad of situations in which the assembly of entire genomes is not the primary objective, but instead the goal is the assembly of one or many discreet, relatively small subgenomic targets. ARC is an iterative algorithm that uses an initial set of reference sequences (subgenomic targets) to seed *de novo* assemblies. Reads are first mapped to reference sequences, and then the mapped reads are pooled and assembled in parallel on a per-target basis to form target-associated contigs. These assembled contigs then serve as reference sequences for the next iteration (see Figure 1). This method breaks the assembly problem down into many smaller problems, using iterative mapping and *de novo* assembly steps to address the poor scaling issue inherent to some *de novo* assembly methods and the reference bias inherent to traditional read mapping. Finally, ARC produces contigs that are annotated to the reference sequence from which they were initiated from, making across sample comparisons possible with little additional processing.

**Figure 1.**
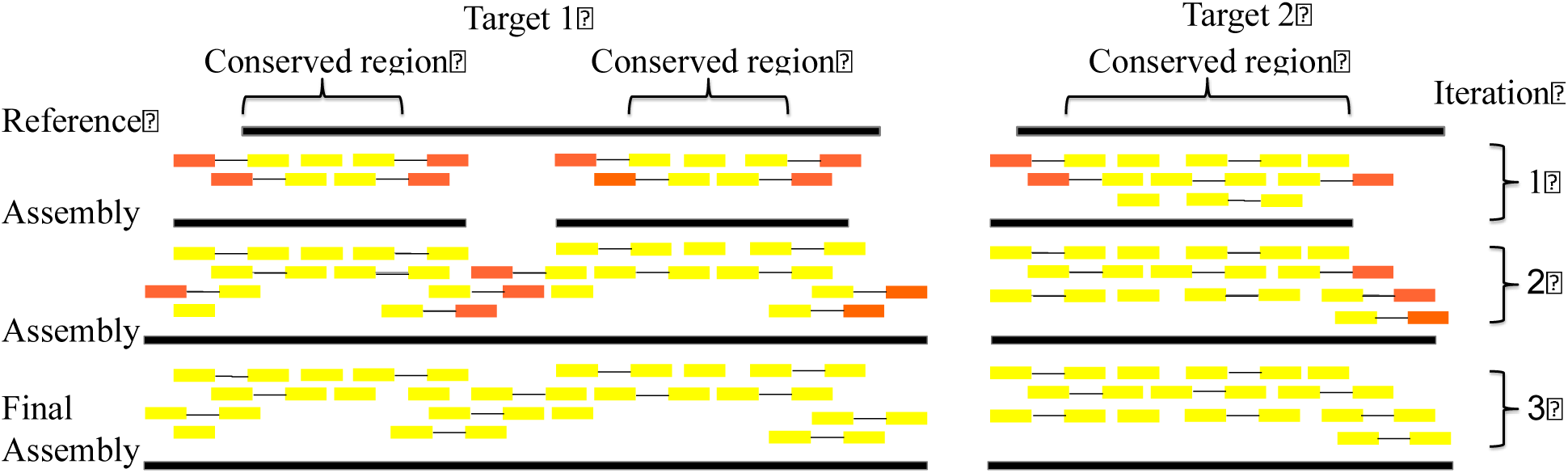
An example of iteratively assembling homologous sequences using ARC. In iteration 1, a small number of reads and unmapped pairs are recruited to the more highly conserved regions of the divergent reference. These reads are assembled and the resulting contigs are used as mapping targets in the next iteration. This process is iterated until no more reads are recruited. Mapped reads are indicated in yellow, unmapped reads in orange. Paired reads are indicated with a connector. Both members of a pair are recruited if only one maps.

## RESULTS

Experiments were conducted to determine how well ARC performs across an array of progressively more divergent references, assembly of short, poor quality reads produced from ancient DNA samples, and to measure ARC’s performance on a large dataset. ARC was tested using two datasets. The first dataset is made up of Illumina sequence reads from two chipmunk (Tamias sp.) exome capture experiments (Bi et al. 2012; Sarver et al. in prep). The second dataset consists of Roche 454 FLX sequence reads from a whole-genome shotgun sequencing experiment using ancient DNA extracted from a mammoth hair shaft sample (Gilbert et al. 2007). The workflow and results of these experiments are presented below.

### Assembly by Reduced Complexity Workflow

The iterative mapping and assembly principle (Figure 1) and workflow (Figure 2) behind ARC consists of several steps: 1) align sequenced reads to reference targets, 2) use alignment results to distribute reads into target specific bins, 3) perform assemblies for each bin (target) to produce contigs, and 4) replace initial reference targets with assembled contigs and iterate the process until stopping criteria have been met. During the read alignment step (1), either the sequence aligner BLAT, or Bowtie 2, is used to identify reads that are similar to the current reference targets. The assembly step (3) is performed using either the Roche GS De Novo Assembler (aka “Newbler”) or SPAdes assemblers.

**Figure 2.**
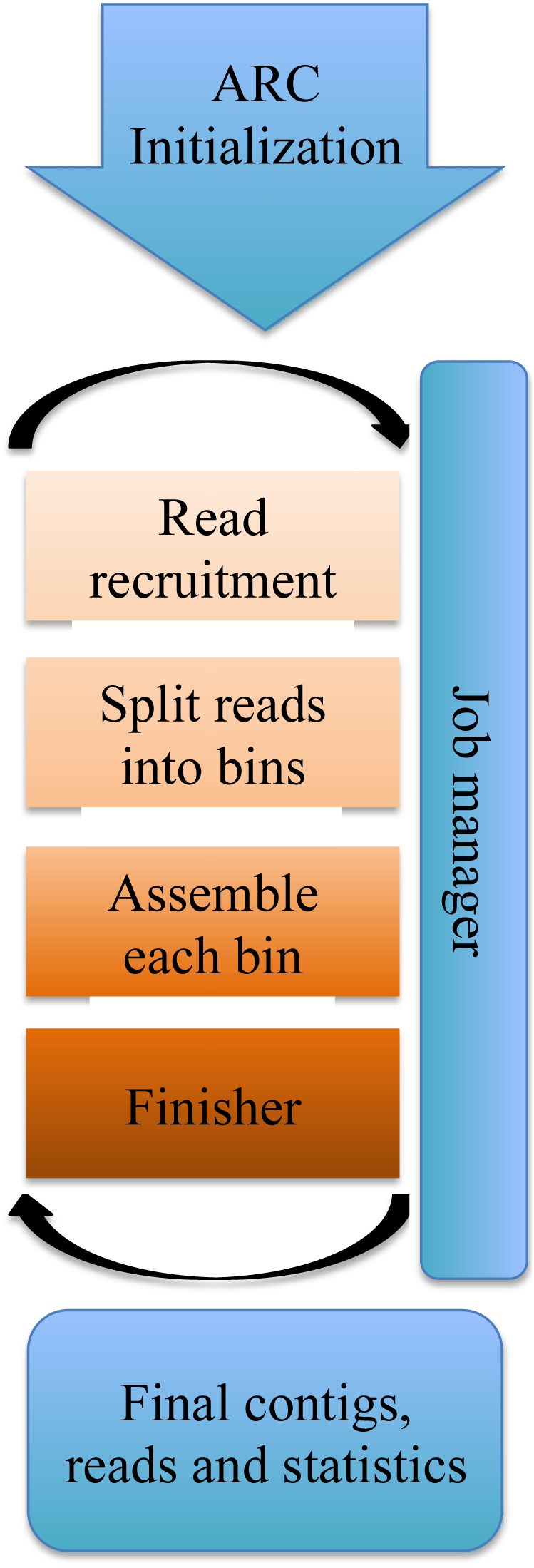
ARC processing stages. The ARC algorithm consists of an initialization stage, followed by four steps: 1) read recruitment, 2) split reads into bins, 3) assemble each bin and 4) finisher. These steps are iterated until stopping conditions are met, at which point a final set of contigs and statistics are produced.

ARC accepts a plain text configuration file, a FASTA formatted file with reference target sequences, and either FASTA or FASTQ formatted files containing reads for each sample. An output folder is generated for each sample that contains the final set of contigs, the reads recruited on the final iteration, and ARC statistics.

ARC is open source software implemented in the Python programming language with source code available for download from GitHub (http://ibest.github.io/ARC/). Prerequisite software packages include: Python 2.7.x, Biopython (Cock et al., 2009), BLAT (Kent, 2002) or Bowtie 2 (Langmead and Salzberg, 2012) and Newbler (Margulies et al., 2005) or SPAdes (Bankevich et al., 2012). These software packages are all free and easy to obtain, and may already be available on systems previously used for HTS analysis. ARC can be installed on most Linux servers, but will also work on many desktops or laptops, provided the required prerequisites are installed. The installation size is only 3Mb, and system administrator access is not required, making it easy to download and use. Configuration is done via a plain text file that can be distributed to make replication of results simple.

### ARC performs well with divergent references

A divergent reference sequence can result in unmapped and misaligned reads (Li, 2012). To test how robust ARC is to reference sequence divergence, we assembled mitochondrial genomes using reads from an exome sequence capture experiment performed on 55 chipmunk specimens representing seven different species within the Tamias genus (*T. canipes*, *T. cinereicollis*, *T. dorsalis*, *T. quadrivittatus*, *T. rufus*, *T. umbrinus*, and *T. striatus*) (Bi et al. 2002; Sarver et al. In prep.). We ran ARC using a set of 11 mitochondrial references spanning Mammalia, including Eastern long fingered.bat, Cape hare, Edible dormouse, Gray-footed chipmunk, Guinea pig, House mouse, Human, Platypus, Red squirrel, Ring-tailed lemur, and Tasmanian devil. Sequence divergence of mitochondrial references with respect to *Tamias cinereicollis* ranged (in percent identity) from 71.2% (Platypus) to 94.9% (Gray-footed chipmunk). Generally speaking, the more divergent the reference sequence, the more ARC iterations were needed in order to complete the assembly process (see Figure 3), while still producing the same resulting mitochondrial genome sequence.

**Figure 3.**
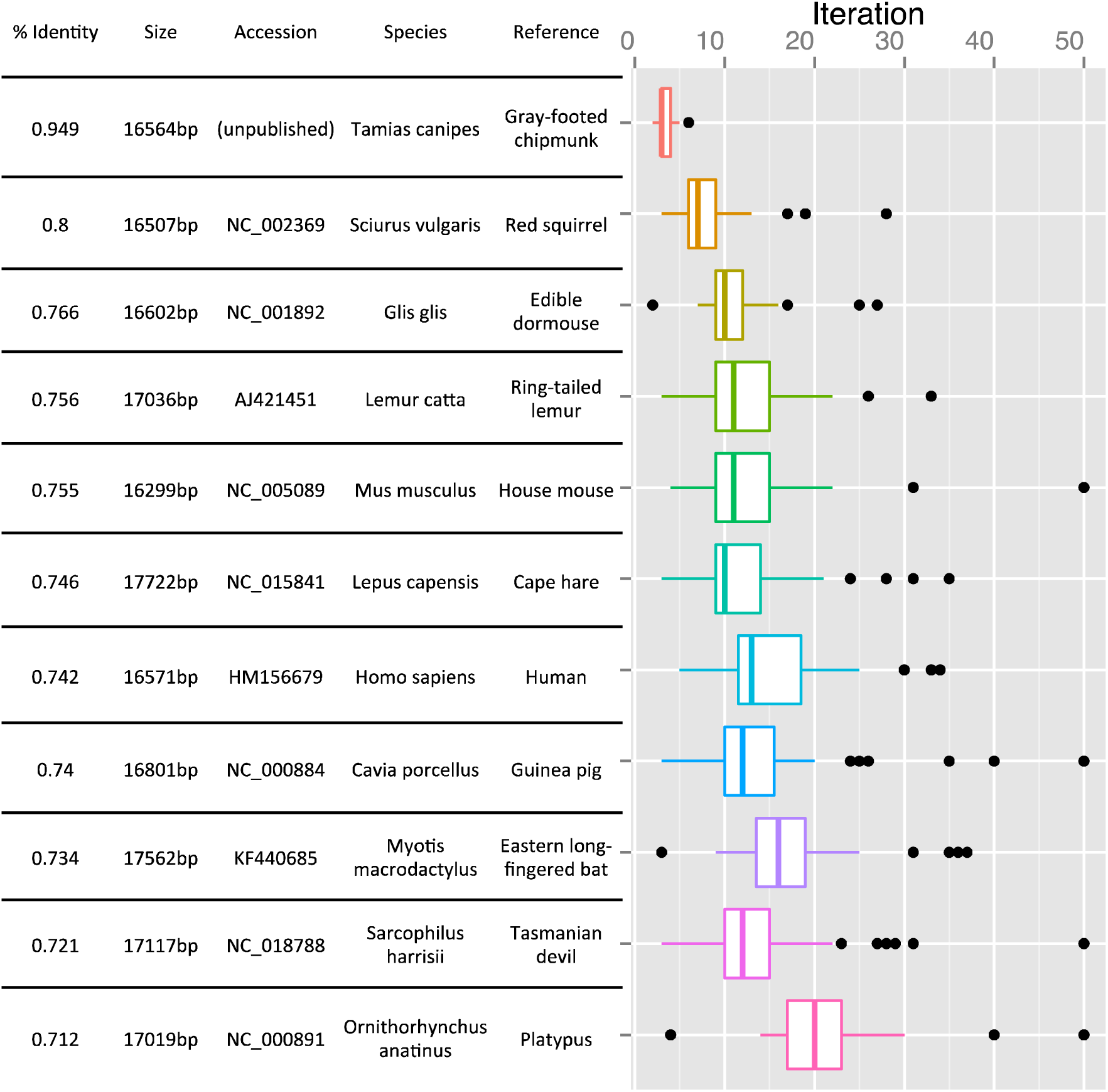
Set of references used for ARC assembly of chipmunk mitochondrial genomes and their respective scientific names, genome sizes, and NCBI Genbank accession numbers. Percent identity is determined with respect to the Gray-Collared chipmunk (*Tamias cinereicollis*). Boxplots show the variation around the number of ARC iterations for each reference species across all 55 samples, before stopping conditions were met.

Supplemental Table 1 reports ARC results for final number of reads recruited and used for assembly (as well as the common count of reads across all 11 reference targets), contig size (total sum of bases across all contigs produced), contig count, ARC iterations needed before stopping criteria were met, and final ARC status (completed or killed) across the 11 reference target sequences for each of the 55 samples. Results show that the choice of reference sequence did not qualitatively impact the final result, ultimately producing, in most cases, the same final mitochondrial genome sequence. In general each of the 11 reference target species recruited the same number of reads, produced the same number of contigs,and resulted in the same length product, with the primary difference being the number of ARC iterations conducted before stopping conditions were met. The relationship between target and read recruitment is further illustrated in Supplemental Figure 1, which shows that the most similar target, the Gray-footed chipmunk (*T. canipes*), typically recruits almost the full set of reads in the first iteration and finishes by the third iteration. At the other extreme, platypus recruited a significantly smaller proportion of reads on the first iteration, but continues to recruit more reads at each iteration until it acquires the full (and often, the same) set of reads. Quality of the original read dataset attributed more to determining the success of assembly than choice of reference sequence.

**Table 1.** ARC results for assembly of ancient mammoth DNA sequences. ARC produces a small number of contigs in all cases with good coverage and identity between the assembled contigs and published reference.

| Reference | Contig count | Total contig length (bp) | Percent coverage | Percent identity | ARC iteration | Reads recruited |
| --- | --- | --- | --- | --- | --- | --- |
| *Mammuthus primigenius* | 4 | 16,620 | 99.7% | 98.1% | 3 | 4633 |
| *Elephas maximus* | 4 | 16,603 | 99.7% | 98.2% | 5 | 4631 |
| *Mus musculus* | 2 | 15,781 | 95.9% | 99.4% | 78 | 4507 |

We observed some variation in final contig lengths across the reference sequences; however, this can be attributed to the linearization of the circular mitochondrial genome. As an example, ARC assembled a single contig for sample S160 across all 11 references, with the length of contig differing by 29 bp between two groups of targets: Edible dormouse, Ring-tailed lemur, and Eastern long-fingered bat targets produced an identical 16,642 bp contig, and all other references produced an identical 16,671 bp contig. A combination of pairwise alignments and dot-plots (data not shown) indicate that these differences are due to the way in which this circular sequence was linearized. The 16,642 bp contig has a 90 bp overlap between the beginning and end of the contig, while the 16,671 bp contig has a 119 bp overlap, caused from group 2 recruiting one additional read relative to group 1. Therefore, even though the assembled length differed slightly the resulting mitochondrial genomes were identical and equal in length after trimming overlapping ends.

### ARC assembles large contigs from short, poor quality reads produced from ancient DNA

Methods that permit investigators to extract DNA from samples that are as much as 50,000 years old and prepare libraries for HTS have been developed (Gilbert et al. 2007, 2008; Knapp and Hofreiter 2010). The DNA from these ancient samples tends to be partially degraded resulting in shorter, poorer quality reads (Knapp and Hofreiter, 2010). As described previously, ARC relies on an iterative process to extend assemblies into gaps. Recruiting reads with partial, overhanging alignments at the edge of a contig eventually fills these gaps. To test the effectiveness of ARC with short, single-end reads produced from ancient samples, we used ARC to assemble the mammoth (*Mammuthus primigenius*) mitochondrial genome using reads sequenced by Gilbert et al. (2007) from DNA collected from hair samples.

Sequenced reads were obtained for *Mammuthus primigenius* specimen M1 from the Sequence Read Archive (SRA001810) and preprocessed as described in the Methods section. ARC was run using three mitochondrial references target sequences: the published sequence from *Mammuthus primigenius* specimen M13, Asian elephant (*Elephas maximus*) the closest extant relative of the mammoth (Gilbert et al. 2008), and a more divergent reference, the house mouse (*Mus musculus*) (accessions: EU153445, AJ428946, NC_005089 respectively).

We evaluated ARC results by alignment to the published *Mammuthus primigenius* M1 sequence (EU153444), which is 16,458 bp in length. Results of this comparison are presented in Table 1. Percent coverage (> 99%) and identity (> 98%) is high for both the mammoth and elephant references. The mouse reference resulted in a slightly smaller assembly (total length 15,781 bp), however coverage (95.9%) and identity (99.4%) were still high. Not surprisingly, the mouse reference required 78 ARC iterations to build its final set of contigs, recruiting only 223 reads on the first iteration. Despite starting from such a small number of initial reads, the final iteration recruited 4,507 reads, almost the same number as the other reference sequences, but from a significantly more divergent reference sequence.

All contigs assembled by ARC could be aligned to the published reference sequence, however the lengths of contigs assembled using the mammoth (16,620 bp) and elephant references (16,603 bp) were both longer than the published sequence length (16,458 bp). To investigate whether this was due to a poor quality assembly on the part of ARC, or an error in the published sequence, we aligned the ARC contigs produced from the mammoth reference (*Mammuthus primigenius* M13) and the published *Mammuthus primigenius* M1 sequence against the published Asian elephant sequence (Supplemental Figure 2). The alignment showed a number of gaps existed in the ARC assembly as compared to the published contigs. Each of these gaps was associated with a homopolymer (consecutive identical bases, e.g., AAA), a known issue with Roche 454 pyrosequencing technology. More interesting was that the D-loop region of the published *Mammuthus primigenius* M1 sequence contains 10 ‘N’ characters followed by a 370bp gap when aligned against the Asian elephant reference. ARC assembled 220 bp of this sequence, including sequence that crosses the unknown, “N” bases in the published sequence. These assembled bases align with high identity against the Asian elephant reference, suggesting that they represent an accurate assembly of this locus and that the published M1 mitochondrial sequence is either missing sequence or is misassembled in this region.

### ARC computational requirements for large datasets

To be useful for modern genomic experiments ARC must be able to process large datasets with multiple samples and potentially thousands of targets. We benchmarked ARC’s performance with the previously described chipmunk exome capture dataset that contains reads from 55 specimens and exonic sequence captured from 7,627 genes as well as the full mitochondrial genome. After stringent read cleaning to remove adapters, PCR duplicates, and overlapping of paired-end reads with short inserts, this dataset contains 21.9 Gbp in 194,597,935 reads. For comparison purposes, we also carried out de novo assemblies of three libraries using the Roche Newbler v2.6 assembler (Table 2).

**Table 2.** ARC assembly of 55 specimens compared to individual R 654 oche Newbler *de novo* assemblies of three specimens (S151, S152, and S223). Maximum and average memory usage (RAM) is listed in gigabytes (GB). Total data processed is reported in millions of base pairs (Mbp)

|  | **ARC** | **Newbler: S151** | **Newbler: S152** | **Newbler: S223** |
| --- | --- | --- | --- | --- |
| Total running time | 77hr, 45min | 31 min | 1hr 13min | 13hr 27min |
| Average Memory (GB) | 22.78 | 5.847 | 8.337 | 16.36 |
| Maximum Memory (GB) | 31.19 | 6.71 | 9.967 | 17.54 |
| Total assemblies performed | 1,300,076 | Not Applicable | Not Applicable | Not Applicable |
| Average assemblies per second | 7.03 | Not Applicable | Not Applicable | Not Applicable |
| Library Size (Mbp) | 21,913 | 243 | 367 | 629 |

ARC required 77 hours 45 minutes to process all 55 samples and 7,627 genes, carrying out a total of 1.3 million assemblies and using a maximum of 31.19 GB of memory. On average this equates to 1 hour 25 minutes per sample. By comparison, individual whole dataset assemblies for a representative three samples were variable, requiring between 6.71 GB and 17.54 GB of memory, with running times of between 31 minutes and 13 hours 27 minutes to complete using Roche Newbler. Although time and memory requirements are smaller for assembly of an individual sample, the total time required to assemble 55 samples in serial would have been be much greater than the time required by ARC to process all samples on a single machine. Likewise, the total memory usage needed to assemble all 55 samples concurrently on a single machine would exceed the memory usage required by ARC on the same machine. Further, assembly algorithms produce anonymous (unannotated) contigs, requiring significant additional processing and analysis before homologous sequences are identified and can be compared between samples. In contrast ARC contigs are annotated to the target from which they were initiated, facilitating across sample comparisons.

Since ARC breaks a large assembly problem into smaller, more manageable pieces, we postulated that memory requirements would scale as a function of the number of CPUs used to perform ARC assemblies rather than as a function of the total number of reads as is normally the case with sequence assembly (Li 2012). To test this, we performed nine ARC runs using between 10 and 50 CPU cores with the 55-specimen chipmunk dataset. We used a random subset of 200 targets instead of the full 7,627 targets so that the experiment could be completed in a reasonable amount of time. During each assembly we recorded maximum memory usage. The results indicate a linear increase in memory usage as the number of cores increases (see Figure 4). A linear model was fit to this data resulting in an estimated slope of 0.07 GB per CPU core (P < .005, R^2^ = 0.96) for this dataset. It is important to note that even though this dataset contains 21.9 Gbp of reads, analysis using a small number of CPU cores and a reduced dataset required less than 3 GB of RAM total, making it possible to use ARC on any size dataset with any modern desktop computer.

**Figure 4.**
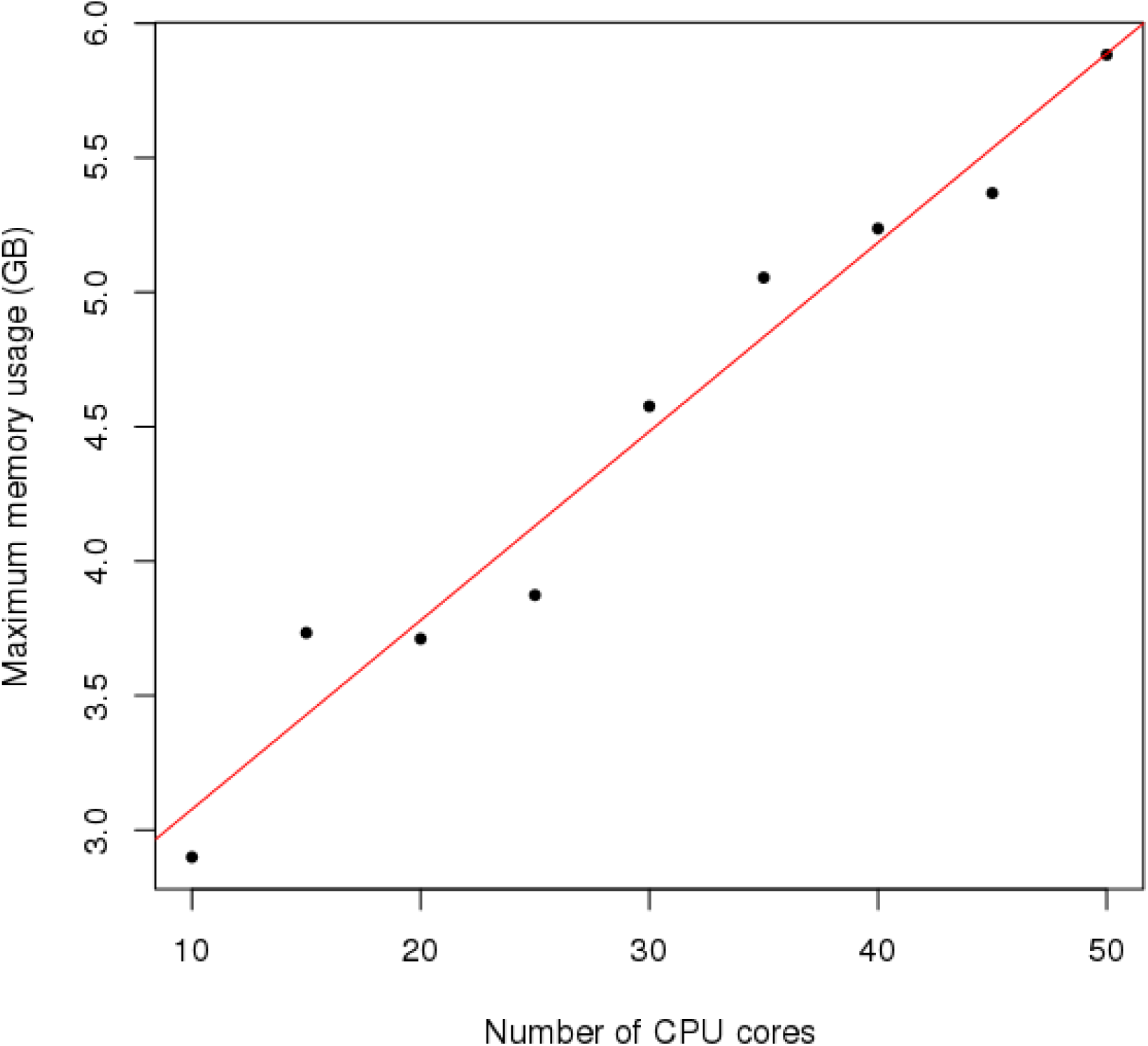
ARC memory requirements (y-axis) scale as function of the number of CPU cores used (x-axis). A line of best fit is plotted in red.

## DISCUSSION

In this paper we introduce ARC, a software package that facilitates targeted assembly of HTS data. This software is designed for use in situations where assembly of one or several discreet and relatively small targets is needed and (potentially divergent) homologous reference sequences are available for seeding these assemblies. ARC fills the gap between fast, mapping based strategies which can fail to map, or misalign reads at divergent loci, and de novo assembly strategies which can be slow, resource intensive, and require significant additional processing after assembly is complete. ARC was evaluated in three ways: 1) we determined whether ARC results were biased by divergence of the reference; 2) we tested the effectiveness of ARC to produce assemblies using short, low quality reads produced from ancient DNA; and 3) we characterized performance on a large HTS dataset with 55 samples and thousands of subgenomic targets.

Assemblies using a divergent set of references with chipmunk specimens show that ARC does not require a close reference to produce high quality final contigs. Supplemental Figure 1 illustrate that on the initial iteration, ARC is able to map only a tiny fraction of the mitochondrial reads to all but the most closely related gray-footed chipmunk reference, yet is able to recover, in most cases, a full set of reads and complete mitochondrial genomes by iteration 50. This small set of reads represents the total number of reads that would have been aligned using a traditional mapping strategy and further illustrates how sensitive read mapping is to high levels of divergence. A similar pattern emerged when we used a mouse reference to seed assembly of a mammoth mitochondrial genome. A mere 223 reads mapped on the first iteration, which was sufficient to seed assembly of an almost full-length mitochondrial sequence assembled from 4,507 reads.

Repetitive sequences and excess coverage are well-known issues, which increase memory usage and slow assembly (Li 2012; Miller et al. 2010). Although ARC partially addresses this problem by breaking the full set of reads into smaller subsets before assembly, it can still encounter issues with very high coverage libraries, or when a target includes repetitive sequence and recruits a large numbers of similar reads. For example, when testing ARC’s ability to handle diverse mitochondrial references, assemblies did not complete for specimen S10 using any of the 11 reference target sequences. In this case the sequence depth was ~1500x for the mitochondrial genome; this depth is not suited for the Newbler assembler, which performs pairwise comparisons of every read and works best when coverage is closer to an expected depth of 60x. The excess coverage led to long assembly times and an eventual timeout. Although the iterative ARC process did not run to completion in this case, intermediate contigs are still reported and contained the full, although fragmented, mitochondrial genome.

ARC has a number of built in mechanisms to mitigate problems caused by repetitive sequences and excess coverage. These include a masking algorithm that inhibits recruitment of reads from simple tandem repeats, as well as tracking of read recruitment patterns that quits assembly if an unexpectedly large number of reads are recruited between iterations, and an assembly timeout parameter that terminates assemblies that run beyond a specified limit. In addition to these strategies there is also an option to subsample reads in cases of known very high sequence depth. Subsampling was not used in any of the tests described here, but may have improved results for samples such as S10. During testing and development, we observed improved behavior with each of these measures on large datasets while minimizing the impact of excess sequencing coverage and repeat elements. Implementing them has allowed ARC to run more quickly and efficiently; however, it is clear that in some cases, recruitment of excess reads and repeat elements can still cause problems for some targets or samples. In all completed assemblies, the resulting set of reads and contigs were either identical or nearly so, providing strong evidence that ARC is able to assemble high quality, unbiased contigs using even very divergent references. This capability makes ARC a very useful tool when analyzing sequence data from non-model organisms or when the identity of a sample is in question.

We tested ARC’s ability to assemble contigs with short, low quality reads recovered from ancient mammoth DNA and found that read length and quality did not impact ARC’s ability to assemble full length genomes. The resulting mitochondrial genome assemblies appear to be as good as or even better than the published assembly for this sample despite using a divergent reference for ARC. Assembly of the M1 mammoth sequence by Gilbert et al. (2007) was achieved through mapping against another mammoth mitochondrial sequence published by Krause et al. (2006) that was generated using a laborious PCR-based strategy. Because ancient DNA sequencing projects are often targeted at extinct organisms (Knapp and Hofreiter 2010) there is rarely a high quality reference from the same species that can be aligned and mapped to. This makes ARC an excellent choice for this type of data, where a target sequence from a related, extant organism is likely to successfully seed assembly. Even in the case where no closely related organism exists, a more distance reference may still be available, as was demonstrated by the assembly of two large contigs representing ~96% of the mammoth mitochondrial genome using a mouse mitochondrial genome for a reference. Additionally, ARC can be configured to use multiple reference sequences as a single target. In cases where specimens cannot be identified, the user can select a set of potentially homologous targets from many phylogenetically diverse taxa so that all sequences may serve as references in order to seed assembly.

Analysis of HTS data can be computationally intensive, and time and memory requirements can become serious limitations, especially with larger datasets (Zhang et al. 2011). With ARC, we have attempted to reduce these requirements using a ‘divide and conquer’ approach that breaks large HTS datasets up into many smaller problems, each of which can be solved quickly and with reduced resources. In the large, 55 sample, 7,627 target dataset, ARC completed over 1.3 million assemblies, averaging seven assemblies per second, in less than 78 hours. This approach allows the user to control memory usage simply by changing the number of CPU cores available to ARC as shown in Figure 4. Less than 3 Gb of RAM was required when using 10 cores, despite processing a 21.9 Gbp dataset that would have required many times this amount of memory using traditional assembly methods. Of course, using fewer CPUs comes with the cost of a longer run time, so ARC can be tuned to the resources available.

It is useful to think of the DNA sequence mapping problem as a trade-off between sensitivity and specificity (Fonseca et al. 2012). To avoid mapping reads to multiple loci throughout the reference, mapping parameters must be tuned for high specificity. However, when divergent loci exist within the reference sequence, high specificity limits the sensitivity of the mapper, leaving reads unmapped. Assembly, on the other hand, can be seen as mapping reads against themselves, thereby removing difficulties associated with divergent reference loci, but incurring the burden of pairwise read comparisons that is significant in large datasets. ARC circumvents these problems by removing reference bias through an iterative mapping and assembly process. As the intermediate reference is improved, more reads can be recruited without sacrificing specificity, allowing both specificity and sensitivity to remain high. At the same time, because only a small subset of reads is assembled, the all-by-all comparisons are less burdensome. This process is carried out in an automated, easily configured manner, with standardized output that simplifies additional analysis, or integration into existing sequence analysis pipelines.

## METHODS

The ARC algorithm proceeds through a number of stages, which have been outlined below and are presented in Figure 1. This algorithm consists of four steps: mapping, splitting, assembling, and finishing. A graphical representation of the algorithm is presented in Figure 1, while an example illustrating the ARC process from the perspective of reads and contigs is provided in Figure 2.

### Initialization

During the initialization stage a configuration file is processed and a number of checks are carried out to ensure that data and prerequisite applications specified in the configuration file are available. If any checks fail, ARC will report an informative error message providing details about the problem and then exit. If all checks pass successfully the initialization process continues by creating internal data structures to store information about the experiment and pipeline progress. Working directories and read index files are created for each sample, and names that are file-system safe are assigned to each reference target sequence. Finally, the job manager is started (including job queues and workers), and read recruitment jobs are added to the job queue for each sample. With initialization complete, ARC begins the iterative part of the pipeline.

### Read recruitment: reads are recruited by mapping against a set of reference target sequences

In the first iterative stage, ARC recruits reads by mapping them against a set of reference targets using one of the two currently supported mappers, BLAT (Kent, 2002) or Bowtie 2 (Langmead and Salzberg, 2012), which is specified in the configuration file. In all subsequent iterations, the reference targets consist of contigs assembled from the previous iteration and are therefore highly similar and no longer represent a divergent reference sequence since they were derived from the sample reads.

BLAT is a fast, seed-and-extend sequence alignment tool that supports gapped alignments and has proven effective at recruiting reads even in cases where global sequence identity is as low as 70%. In the first iteration, BLAT is run using default parameters (minIdentity=90, minScore=30) but on all subsequent iterations mapping stringency is increased (minIdentity=98, minScore=40) to reduce recruitment of less similar reads. BLAT reports all alignments that meet the minimum score criteria, so it is possible to use the same read multiple times if it aligns successfully against more than one target. One drawback of using BLAT is that it does not support the FASTQ format. All current HTS platforms produce base quality information for reads and this information is typically encoded in FASTQ format. To facilitate usage of ARC and FASTQ formatted data we include a code patch for BLAT that adds support for FASTQ files. Instructions for applying this patch can be found in the online manual http://ibest.github.io/ARC/).

Bowtie 2 is another fast, gapped, read aligner that was specifically designed for mapping HTS reads (Langmead and Salzberg, 2012). Bowtie 2 is ran in ARC under local alignment mode (--local option) which enables the recruitment of reads that partially map to the ends of contigs and in low-homology regions. Additionally, the option to report up to five valid alignments (-k 5) is used by default. This setting can be modified based on the user’s expectations by setting the bowtie2_k parameter in the ARC configuration file. Setting bowtie2_k=1 will cause Bowtie 2 to run in default local-alignment mode where only the best alignment found is reported.

### Split reads into bins: reads are split into subsets based on mapping results

In the second iterative stage, ARC splits reads into bins based on the mapping results. The supported mappers, BLAT and Bowtie 2 generate PSL or SAM (Li et al. 2009) formatted output files, respectively. ARC processes each sample’s mapping output file and reads are split by reference target. This is accomplished by creating a series of FASTQ files corresponding to reads which map to each reference target; allowing for the assembly of each target’s reads independently from the others. Splitting requires fast random access to the read files, which is facilitated by storing read offset values in a SQLite database as implemented in the Biopython SeqIO module (Cock et al. 2009). Two special considerations are taken into account during splitting. First, since the Newbler assembler uses pre CASAVA 1.8 Illumina read identifiers to associate paired reads, it is necessary to reformat the read identifier to ensure compatibility with Newbler paired-end detection. This is performed by ensuring that the read identifier is made up of five fields separated by a colon and ending in a sixth field indicating the pair number, a format compatible with most modern day assemblers. Identifiers for single-end reads are similarly reformatted, except that the sixth field, which indicates pair number, is left blank. Secondly, regardless of whether one or both of a read pair map to a target, both members of the pair are recruited as long as at least one of them was mapped to the target sequence. Recruiting paired reads in this way takes advantage of the information stored in paired reads, and allows for faster extension of targets.

Despite using a fast strategy for random accessing of read files, splitting is limited by system input/output latency and to a single CPU core per sample. To optimize CPU use on modern multi-core systems, ARC immediately adds an assembly job to the job queue as soon as all reads associated with a target have been split. This allows assemblies to proceed concurrently with the read splitting process.

### Assemble each bin: targets are assembled using either the Spades or Newbler assemblers

Because the read splitting process is carried out sequentially across mapping reference targets, an assembly job for a target can be launched as soon as all reads associated with the target have been written. As soon as resources become available, assembly jobs are started, allowing ARC to run read splitting and assembly processes concurrently. Two assemblers are currently supported, the Roche GS *de novo* Assembler (also known as Newbler; Margulies et al., 2005), and SPAdes (Bankevich et al., 2012). Assemblies within ARC are always run with a timeout in order to gracefully handle the cases where the assembler crashes, does not exit properly, or takes longer than expected to run. This allows ARC to continue running efficiently on large projects where a small number of targets might be problematic (e.g., due to recruiting reads from repetitive elements). The timeout value can be controlled using the assembly timeout setting in the configuration file.

Newbler was originally designed to assemble reads generated from the Roche 454 pyrosequencing platform (Margulies et al. 2005), but recent versions have added support for Illumina paired-end reads and Newber can be run using only Illumina reads. The ARC configuration file supports two Newbler specific parameters that can sometimes improve assembly performance. These are to set urt=True, which instructs Newbler to “use read tips” in assemblies, and rip=True, which instructs Newbler to place reads in only one contig and to not to break and assign reads across multiple contigs. We have found that setting urt=True can reduce the number of ARC iterations necessary to assemble a target.

The second assembler supported in ARC is SPAdes (Bankevich et al., 2012). SPAdes is an easy to use de Bruijn graph assembler that performed well in a recent evaluation of bacterial genome assemblers (Magoc et al., 2013). SPAdes performs well in the ARC pipeline, but is not as fast as Newbler for small target read sets (data not shown). This may partly be because SPAdes implements a number of steps in an attempt at improving the often-fragmented de Bruijn graph assembly results seen in large eukaryotic genomes. These steps include: read error correction, multiple assemblies using different k-mer sizes, and merging of these assemblies. In ARC, SPAdes is run using the default set of parameters.

In some cases, the reference targets may be very divergent from the sequenced specimen and, therefore, only a small number of reads are recruited in the first iteration. If too few overlapping reads are recruited, the assemblers have very little data to work with, and in the case of SPAdes, may fail to assemble any contigs. In an attempt to address this specific situation, we provide a final pseudo-assembly option that skips assembly on the first iteration and treats any recruited reads as contigs. These reads are then used as mapping reference targets in the second iteration. This option can be enabled by setting map_against_reads=True in the ARC configuration file. In some cases using reads as mapping targets results in recruiting large numbers of reads from repeat regions, causing the assembly to timeout and fail. For this reason we only recommend using this approach after testing ARC with standard settings.

### Finisher: assembled contigs are written as a new set of mapping targets or to finished output

Once all assemblies are completed for a given sample, the final iterative stage in the ARC pipeline is initiated. During this stage each target is evaluated; if stopping conditions are met, the contigs are written to the final output file; and if not the contigs are written to a temporary file where they are used as reference targets in the next iteration (see the section Folder structure: outputs and logging for details). Stopping conditions within ARC are defined as follows: 1) iterations have reached their maximum allowable number as defined by the numcycle parameter in the ARC configuration file; 2) no additional reads have been recruited (i.e., delta read count between iterations is zero); 3) detection of an assembly that was halted, or killed will result in no further attempts at assembling this target, and any contigs produced on the previous iteration will be written to the output file; or 4) a sudden spike in read counts. Occasionally a target will be flanked by repeated sequence in the genome that can cause a sudden spike in the number of recruited reads. The max_incorporation parameter in the ARC configuration file controls sensitivity to this situation and by default is triggered if five times the previous number of reads are recruited.

During output, target contig identifiers are modified to reflect their sample, original reference target, and contig number separated by the delimeter “_:_” (e.g. sample_:_original-reference-target_:_contig). Contigs are also masked of simple tandem repeats in all but the final iteration, using an approach that relies on frequency of trinucleotides in a sliding window. Repeats are masked by setting them to lower case for Blat support, or by modifying the repeat sequence to the IUPAC character ‘N’ for Bowtie 2 support. All target contigs in their final iteration are written to the final output file, and all corresponding reads are written to the final read files, however their description field is modified to reflect which reference target they are assigned to.

For any targets that remain unfinished (i.e., stopping conditions have not been met), those reference targets are iterated using the newly assembled contigs as the next mapping reference targets.

### Description of input files

Inputs to ARC consist of three types of files: a file containing reference target sequence(s), file(s) containing sequence reads for each sample, and an ARC configuration file.

The reference target sequence(s) file contains the sequences that are to be used as mapping references during the first iteration of ARC. This file must be in standard FASTA format and should have informative, unique names. It is possible to use multiple reference sequences as a single target in cases where a number of potentially homologous targets are available and it is not clear which of them is most similar to the sequenced sample (e.g. in the case of ancient DNA extracted from unidentified bone material). This can be accomplished by naming each reference target using ARC’s internal identifier naming scheme made of three parts separated by “_:_” (e.g., sample_:_reference-target_:_contig). During read splitting, ARC will treat all sequences that have an identical value in “reference-target” as a single target.

Sample sequence read files are represented with up to three sequence read files; two paired-end (PE) files, and one single-end (SE) file. ARC will function with only one SE file, a PE set of files, or all three files if provided. If multiple sets of reads are available for a single biological sample (i.e., from different sequencing runs or technologies) they should be combined into the above described three read files. All reads for all samples must be in the same format (i.e., FASTA or FASTQ) and this format needs to be indicated using the format parameter in the ARC configuration file. It is highly recommended that reads be preprocessed to remove adapter sequences and low quality bases prior to running ARC. Removing PCR duplicate reads and merging paired-end reads has also been observed to produce higher quality, less fragmented ARC assemblies, particularly with capture data (data not shown).

The ARC configuration file is a plain text file describing the various parameters that ARC will use during assembly, mapping, and output stages and the sample(s) read data data paths. By default the configuration file should be named ARC_config.txt, but any name can be used as long as the -c filename switch is used. The configuration file is split into three parts, denoted by the first characters in the line. Lines starting with the characters “##” are treated as comments and ignored, lines starting with “#” are used to set ARC parameters, and lines that don’t begin with “#” indicate sample read data. The one exception to this rule is the sample read data column header line, which is the first line that doesn’t begin with “#”, and contains column names. This line is ignored by ARC, but is expected in the configuration file. An example ARC configuration file is included in the “test_data” directory that comes with ARC. A comprehensive list of configuration options are presented in the online manual (http://ibest.github.io/ARC/).

### Folder Structure Outputs and Logging

In order to minimize memory usage and interface with assembly and mapping applications, ARC relies heavily on temporary files. These files are organized into subdirectories under the path from which ARC is launched. During ARC processing a pair of folders is created for each sample. These folders have the prefixes “working_” and “finished_”. Temporary files used during ARC processing are stored in the “working_” folders while completed results and statistics are recorded in the “finished_” folders.

The “working_” directories contain the sample contigs assembled during each iteration in a set of files with file names “I00N_contigs.fasta” (where “N” corresponds to the iteration) and the latest assembly directory denoted by “t__0000N” (where “N” corresponds to the numeric index of the target). These directories and files can be informative in determining why an assembly failed or for examining assembly statistics of a particular sample and target in more depth. Additionally, these folders provide the option of manually re-running an assembly with a different set of parameters than those chosen within ARC. In addition to the per iteration contigs and latest assembly directories, the “working_” folders also contain the sample read indexes, which can be reused when re-running ARC with new parameters, and the latest mapping log report. The “working_” folders only contain temporary files used by ARC and can be safely deleted after the ARC run.

The “finished_” directories contain the following files: contigs.fasta, mapping_stats.tsv, target_ summary_table.tsv, and final read files. The contigs.fasta file contains the final set of assembled contigs for each target. Contigs are named according to the three part naming scheme previously described (sample_:_original-reference-target_:_contig) in order to facilitate downstream comparisons between samples. The mapping_stats.tsv and target_summary_table.tsv files are tab-separated values files that store information on the number of reads mapped to each target at each iteration and per target final summary statistics respectively. These files can be easily loaded into a spreadsheet, or statistical program such as R to generate plots or for other downstream analysis. The final read files (PE1.fasta/PE1.fastq, PE2.fasta/PE2.fastq, and SE.fasta/SE.fastq) contain all the reads that were mapped, and consequently used during assembly, on the final ARC iteration. If only pair-end or single-end files were provided then only reads of this type will be reported. These files will be formatted in the same manner as the original input files (FASTA or FASTQ) and have modified description fields to indicate the sample and target to which they were assigned.

### ARC post processing and contrib scripts

ARC contains a number of add on scripts in the “contrib” folder of the application, for downstream processing of assembled contigs and visualization of ARC results. These scripts include R functions to profile and plot memory usage and to plot data from the run log. The contrib folder also contains number of Python scripts for post-processing ARC contigs for use in downstream applications such as phylogenomics. Two scripts in particular are “ARC_Add_Cigar_Strings.py” and “ARC_Call_and_Inject_hets.py”. The first allows users to determine the order and orientation of ARC-generated contigs relative to the original reference, using the program BLAT to align assembled contigs against sequences from the original reference targets sequence file. The script then generates a CIGAR string in standard SAM format to describe the alignment. In situations where the contig extends beyond the 5’ or 3’ ends of the target sequence, those bases are described as soft-clipped. The order of the CIGAR string depends on the orientation of the contig with respect to the target (as is the case with similar programs such as Bowtie2). If the contig maps to the forward strand, the CIGAR string reports the matches, insertions, deletions, and soft-clipped regions of the alignment in the 5’ to 3’ direction. In contrast, if the contig maps to the reverse strand, the CIGAR string reports components of the alignment in the 3’ to 5’ direction. The script generates an output file (in FASTA format) that includes the contig sequence from the original ARC output file, the name of the contig, the name of the target sequence the contig mapped to, the start and end positions of the contig relative to the target sequence, the contig’s orientation (i.e., “+” or “-” depending on whether the contig mapped to the forward or reverse strand of the target), and the CIGAR string. With this information the user can ascertain the order and orientation of ARC-generated contigs with respect to the reference.

The second script, “ARC_Call_and_Inject_hets.py”, produces both a variant call formatted file (VCF) per sample and a new contigs file with ambiguity bases at heterozygous loci. This script uses Bowtie 2 to map the reads recruited for each target to their respective assembled contigs. GATK and Picard Tools are then used to call heterozygous SNPs and output a VCF file for each sample. Finally, the script encodes the heterozygous SNP calls using their respective IUPAC ambiguity code and ‘injects’ those bases into the original contig sequences producing a new contigs file containing heterozygous sites.

### Datasets used for testing

We tested ARC with two datasets. The first dataset is made up of Illumina sequence reads from two chipmunk (Tamias sp.) exome capture experiments. This combined dataset consists of sequence reads from 55 specimens, 3 of which were sequenced as part of Bi et al. (2012) while the other 52 were sequenced as part of a separate study (Sarver et al. in prep). The second dataset consists of Roche 454 FLX sequence reads from a whole-genome shotgun sequencing experiment using ancient DNA extracted from a mammoth hair shaft sample (Gilbert et al. 2007).

The first chipmunk dataset was used to investigate ARC’s sensitivity to divergent references as well as its utility and performance with large datasets. For all 55 specimens, libraries were captured using an Agilent SureSelect custom 1M-feature microarray capture platform that contains 13,000 capture regions representing the mitochondrial genome and 9,716 genes (Bi et al. 2012). Libraries were then sequenced on the Illumina HiSeq 2000 platform (100bp paired-end). The 55 chipmunks represent seven different species within the genus Tamias with representatives of T.canipes: 5, T. cinereicollis: 9, T. dorsalis: 12, T. quadrivittatus: 1, T. rufus: 5, and T. umbrinus: 10, collected and sequenced as part of Sarver et al. (in prep) and T. striatus: 3 collected and sequenced by Bi et al. (2012).

Prior to ARC analysis, reads were preprocessed through a read cleaning pipeline consisting of the following steps. PCR duplicates were first removed using a custom Python script. Sequences were then cleaned to remove sequencing adapters and low quality bases using the software package Seqyclean (Zhbannikov et al. in prep, <u>https://bitbucket.org/izhbannikov/seqyclean)</u>. Finally, because paired-end sequencing produces two reads sequenced from either end of a single template, it is often possible to overlap these reads to form a single long read representing the template in its entirety. This overlapping was carried out using the Flash software package (Magoc and Salzberg, 2011). Post-cleaning, the dataset consisted of 21.9 Gbp (giga base pairs) in 194,597,935 reads.

ARC analysis for the first dataset was carried out using two different sets of references. To determine how well ARC performs with divergent references, the mitochondrial genome of each specimen was assembled against eleven different mammalian mitochondrial references (see Figure 3). We also tested ARC’s performance with a large number of targets by using a target set consisting of a manually assembled *Tamias canipes* mitochondrial sequence plus 11,976 exon sequences comprising 7,627 genes. These sequences represent the unambiguous subset from the 9,716 genes that the capture probes were originally designed against.

The second woolly mammoth dataset was used to test ARC’s performance on shorter, poor quality reads that are typical of ancient DNA sequencing projects. Total DNA was extracted from ancient hair shafts and reads were sequenced on the Roche 454 FLX platform by (Gilbert et al. 2007). Although these reads represent shotgun sequencing of both the nuclear and mitochondrial genomes, the authors report a high concentration of mitochondria in hair shaft samples resulting in high levels of mitochondrial reads relative to nuclear reads. Sequenced reads for *Mammuthus primigenius* specimen M1 were obtained from the Short Read Archive using accession SRX001889 and cleaned with SeqyClean (Zhbannikov et al. in prep, <u>https://bitbucket.org/izhbannikov/seqyclean)</u> to remove 454 sequencing adapters and low quality bases. Following cleaning, this datasets contains a total of 19 Mbp in 221,688 reads with an average length of 86.2 bp. Although these reads were sequenced on the Roche 454 platform which typically produces much longer reads (400-700bp), 75% of cleaned reads were 101bp or less in length making them extremely short for this platform. ARC analysis was carried out using three mitochondrial references, the published *Mammuthus primigenius* sequence from another specimen, M13, Asian elephant (*Elephas maximus*) the closest extant relative of the mammoth (Gilbert et al. 2008), and a divergent reference, mouse (*Mus musculus*) (accessions: EU153445, AJ428946, NC_005089 respectively).

## DATA ACCESS

The raw data used in this study are available in NCBI Sequence Read Archives under BioProject numbers SRX001889 (Mammuthus primigenius M1), SRA053502 (Tamias samples S10, S11, S12), and SRAXXXXX (Remaining Tamias samples). Reference sequences used in this study are available in NCBI Genbank under accession numbers: NC_000884.1 (guinea pig), NC_001892.1 (edible dormouse), HM156679.1 (human), AJ421451.1, (ring-tailed lemur), NC_015841.1 (cape hare), KF440685.1 (eastern long-fingured bat), NC_000891.1 (platypus), NC_018788.1 (tasmanian devil), NC_002369 (red squirrel), NC_005089 (house mouse), EU153445 (*Mammuthus primigenius*), AJ428946 (*Elephas maxiumus*), NC_005089 (*Mus musculus*).

## ACKNOWLEDGEMENTS

We would like to thank Ilya Zhbannikov for generating the FASTQ patch to BLAT. We would also like to thank Jeffery Good, John Demboski, Jack Sullivan, Dan Vanderpool, and Kayce Bell for aid in collecting and sequencing chipmunk samples. Research reported in this publication was supported by the National Institute Of General Medical Sciences of the National Institutes of Health under Award Number P30 GM103324 and the National Science Foundation under Cooperative Agreement No. DBI-0939454 and DEB-0717426. Any opinions, findings, and conclusions or recommendations expressed in this material are those of the author(s) and do not necessarily reflect the views of the National Science Foundation.

## REFERENCES

Bankevich, A., Nurk, S., Antipov, D., Gurevich, A. A., Dvorkin, M., Kulikov, A. S., Lesin, V. M., Nikolenko, S. I., Pham, S., Prjibelski, A. D., et al. (2012). SPAdes: A New Genome Assembly Algorithm and Its Applications to Single-Cell Sequencing. Journal of Computational Biology, 19(5), 455–477. doi: 10.1089/cmb.2012.0021

Bi, K., Vanderpool, D., Singhal, S., Linderoth, T., Moritz, C., Good, J. M. (2012). Transcriptome-based exon capture enables highly cost-effective comparative genomic data collection at moderate evolutionary scales. BMC Genomics, 13(1), 403. doi:10.1186/1471-2164-13-403

Bradnam, K. R., Fass, J. N., Alexandrov, A., Baranay, P., Bechner, M., Birol, I., Boisvert, S., Chapman, J. A., Chapuis, G., Chikhi, R., et al. (2013). Assemblathon 2: evaluating de novo methods of genome assembly in three vertebrate species. GigaScience, 2(1), 10. doi: 10.1186/2047-217X-2-10

Chevreux, B., Wetter, T., Suhai, S. (1999). Genome Sequence Assembly Using Trace Signals and Additional Sequence Information. Computer Science and Biology: Proceedings of the German Conference on Bioinformatics (GCB), 45–56.

Cock, P. J. A., Antao, T., Chang, J. T., Chapman, B. A., Cox, C. J., Dalke, A., Friedberg, I., Hamelryck, T., Kauff, F., Wilczynski, B., et al. (2009). Biopython: Freely available Python tools for computational molecular biology and bioinformatics. Bioinformatics, 25(11), 1422–1423.

Fonseca, N. A., Rung, J., Brazma, A., Marioni, J. C. (2012). Tools for mapping high-throughput sequencing data. Bioinformatics.

Gilbert, M. T. P., Tomsho, L. P., Rendulic, S., Packard, M., Drautz, D. I., Sher, A., Tikhonov, A., Dalén, L., Kuznetsova, T., Kosintsev, P., et al. (2007). Whole-genome shotgun sequencing of mitochondria from ancient hair shafts. Science (New York, N.Y.), 317(5846), 1927–1930.

Gilbert, M. T. P., Drautz, D. I., Lesk, A. M., Ho, S. Y. W., Qi, J., Ratan, A., Hsu, C., Sher, A., Dalén, L., Götherström, A., et al. (2008). Intraspecific phylogenetic analysis of Siberian woolly mammoths using complete mitochondrial genomes. Proceedings of the National Academy of Sciences of the United States of America, 105(24), 8327–8332.

Hahn, C., Bachmann, L., Chevreux, B. (2013). Reconstructing mitochondrial genomes directly from genomic next-generation sequencing reads--a baiting and iterative mapping approach. Nucleic Acids Research, 41(13), e129. doi:10.1093/nar/gkt371

Kent, W. J. (2002). BLAT - The BLAST-like alignment tool. Genome Research, 12(4), 656–664.

Knapp, M., Hofreiter, M. (2010). Next Generation Sequencing of Ancient DNA: Requirements, Strategies and Perspectives. Genes, 1(2), 227–243. doi:10.3390/genes1020227

Krause, J., Dear, P. H., Pollack, J. L., Slatkin, M., Spriggs, H., Barnes, I., Lister, A. M., Ebersberger, I., Pääbo, S., Hofreiter, M. (2006). Multiplex amplification of the mammoth mitochondrial genome and the evolution of Elephantidae. Nature, 439(7077), 724–727. doi:10.1038/nature04432

Langmead, B., Salzberg, S. L. (2012). Fast gapped-read alignment with Bowtie 2. Nature Methods, 9(4), 357–359. doi:10.1038/nmeth.1923

Li, H., Handsaker, B., Wysoker, A., Fennell, T., Ruan, J., Homer, N., Marth, G., Abecasis, G., Durbin, R. (2009). The Sequence Alignment/Map format and SAMtools. Bioinformatics, 25(16), 2078–2079.

Li, H. (2011). Improving SNP discovery by base alignment quality. Bioinformatics, 27(8), 1157–1158.

Li, H. (2012). Exploring single-sample snp and indel calling with whole-genome de novo assembly. Bioinformatics, 28(14), 1838–1844.

Magoc, T., Salzberg, S. L. (2011). FLASH: Fast length adjustment of short reads to improve genome assemblies. Bioinformatics, 27(21), 2957–2963.

Magoc, T., Pabinger, S., Canzar, S., Liu, X., Su, Q., Puiu, D., Tallon, L. J., Salzberg, S. L. (2013). GAGE-B: An evaluation of genome assemblers for bacterial organisms. Bioinformatics, 29(14), 1718–1725.

Malé, P. J. G., Bardon, L., Besnard, G., Coissac, E., Delsuc, F., Engel, J., Lhuillier, E., Scotti-Saintagne, C., Tinaut, A., Chave, J. (2014). Genome skimming by shotgun sequencing helps resolve the phylogeny of a pantropical tree family. Molecular Ecology Resources, 14(5), 966–975.

Margulies, M., Egholm, M., Altman, W. E., Attiya, S., Bader, J. S., Bemben, L. A., Berka, J., Braverman, M. S., Chen, Y., Chen, Z., et al. (2005). Genome sequencing in microfabricated high-density picolitre reactors. Nature, 437(7057), 376–380.

Miller, J. R., Koren, S., Sutton, G. (2010). Assembly algorithms for next-generation sequencing data. Genomics.

Picardi, E., Pesole, G. (2012). Mitochondrial genomes gleaned from human whole-exome sequencing. Nature Methods, 9(6), 523–4. doi:10.1038/nmeth.2029

Pyrkosz, A. B., Cheng, H., Brown, C. T. (2013). RNA-Seq Mapping Errors When Using Incomplete Reference Transcriptomes of Vertebrates. arXiv: 1303.2411, 1–17. Retrieved from http://arxiv.org/abs/1303.2411

Sarver

Schbath, S., Martin, V., Zytnicki, M., Fayolle, J., Loux, V., Gibrat, J. F. (2012). Mapping Reads on a Genomic Sequence: An Algorithmic Overview and a Practical Comparative Analysis. Journal of Computational Biology, 19(6), 796–813. doi: 10.1089/cmb.2012.0022

The 1000 Genomes Project Consortium, Abecasis, G. R., Altshuler, D., Auton, A., Brooks, L. D., Durbin, R. M., Gibbs, R. A., Hurles, M. E., McVean, G. A. (2010). A map of human genome variation from population-scale sequencing. Nature, 467(7319), 1061–73. doi:10.1038/nature09534

Zhang, W., Chen, J., Yang, Y., Tang, Y., Shang, J., Shen, B. (2011). A practical comparison of De Novo genome assembly software tools for next-generation sequencing technologies. PLoS ONE, 6(3).

Zhbannikov, I. Y., Hunter, S. S., Foster, J. A., Settles, M. L. (2014) SeqyClean: a pipeline for high throughput sequence data preprocessing. In Prep.

